# Aβ42-Driven α-synuclein Fibril Polymorphism and Distinct Intracellular Aggregation

**DOI:** 10.64898/2026.09.09.750398

**Authors:** Purnopama Saha, Teshani Liyanaarachchige, Anika Alim, Jungwook Paek, Myungwoon Lee

**Author notes:** Corresponding Author: Jungwook Paek, Myungwoon Lee. P.S and T.L contributed equally to this work.

## Abstract

The frequent coexistence of α-synuclein (α-syn) and amyloid-β (Aβ) aggregates in neurodegenerative diseases suggests that heterotypic interactions between these amyloidogenic proteins may influence disease progression, yet their molecular consequences remain poorly understood. Here, we investigated how distinct aggregation states of Aβ42, monomers and preformed fibrils (PFFs), modulate α-syn fibril formation, structure, and downstream neuronal pathology. Thioflavin T kinetics showed that Aβ42 monomers delayed α-syn fibril formation, whereas Aβ42 PFFs exhibited a trend toward accelerated aggregation, indicating aggregation state-dependent effects on α-syn aggregation. Negative-stain TEM, proteinase K digestion, and solid-state NMR spectroscopy further demonstrated that both Aβ42 monomers and PFFs altered α-syn fibril structure, generating distinct fibril conformations depending on the Aβ42 concentration and aggregation state. To determine whether these conformational differences influence pathological activity, α-syn PFF variants generated in the presence of different concentrations of Aβ42 monomers or PFFs were applied to dopaminergic neuronal cells. α-Syn fibrils formed in the presence of Aβ42 PFFs showed greater capacity to induce intraneuronal α-syn aggregation than α-syn PFFs, whereas fibrils formed in the presence of Aβ42 monomers exhibited similar or reduced intraneuronal aggregation-inducing capacity relative to α-syn PFFs. Together, our findings demonstrate that heterotypic interactions with Aβ42 reshape α-syn aggregation pathways and fibril conformations, generating structurally distinct α-syn fibril populations that vary in their ability to trigger intraneuronal aggregation. These results provide a molecular framework for understanding how cross-talk between amyloidogenic proteins may contribute to structural and pathological heterogeneity in mixed neurodegenerative diseases.

## Introduction

Neurodegenerative diseases are commonly characterized by the aberrant deposition of specific protein aggregates, known as amyloid fibrils. For example, synucleinopathies, including Parkinson’s disease (PD), Lewy body dementia (LBD), and multiple system atrophy (MSA), are defined by the accumulation of Lewy bodies (LBs), which are predominantly composed of aggregated α-synuclein (α-syn).^1–6^ In contrast, Alzheimer’s disease (AD), the most prevalent neurodegenerative disorder and a major form of tauopathy, is characterized by extracellular amyloid plaques composed primarily of amyloid-β (Aβ) and intracellular neurofibrillary tau tangles.^7–10^ Accordingly, extensive studies have primarily focused on elucidating the molecular mechanisms and structural properties of aggregates formed by individual amyloidogenic proteins in isolation, yielding important insights into these homotypic fibrils.^11–19^

However, accumulating evidence has revealed that aggregates formed by different amyloidogenic proteins frequently coexist within the same brain regions of affected individuals, suggesting that their aggregation pathways may intersect. For example, Aβ plaques and α-syn aggregates have been observed simultaneously in the cortex of AD patients,^20^ and LBs are detected in the amygdala of AD patients.^21^ Similarly, substantial Aβ deposition is commonly observed in patients with Parkinson’s disease dementia (PDD).^22,23^ This convergence is not merely coincidental: although α-syn and Aβ are preferentially localized to different cellular compartments under physiological conditions, both proteins can access overlapping extracellular and intracellular environments. Extracellular Aβ can be internalized by neurons, whereas α-syn can be released from cells into the extracellular space and subsequently taken up by neighboring cells.^24–26^ These dynamic trafficking processes create opportunities for direct heterotypic interactions between α-syn and Aβ. Together, the co-occurrence of their aggregates at the tissue level and their potential colocalization within shared cellular environments strongly support molecular cross-talk between α-syn and Aβ and raise the possibility that such interactions contribute to co-aggregation and the overlapping pathological features seen across neurodegenerative diseases.

Critically, this molecular cross-talk appears to have direct clinical consequences.^23,27–29^ Patients with mixed Aβ/α-syn pathology often show more rapid cognitive decline, increased mortality, and greater neuropathological burden than those with isolated pathology.^30–33^ Notably, the presence of Aβ plaques has been associated not only with increased cortical α-syn aggregation but also with enhanced spreading of α-syn pathology in PDD and dementia with Lewy bodies (DLB). This observation suggests that Aβ may actively promote α-syn pathology rather than simply co-existing alongside it.^21^ These clinical observations imply that Aβ-α-syn interactions are an active driver of pathological severity, underscoring the need to understand the molecular basis of this interaction directly.

Experimental studies further substantiated a functional interaction between α-syn and Aβ.^34,35^ In animal models, injection of α-syn preformed fibrils (PFFs) markedly enhances Aβ deposition throughout the brain.^36,37^ Conversely, pre-existing Aβ plaques promote α-syn aggregation in vivo, leading to exacerbated neuroinflammation in mice harboring both α-syn and Aβ pathologies.^38,39^ Consistent with these findings, doubly transgenic mice co-expressing α-syn and Aβ precursor protein exhibit severe neurological deficits than either single-transgenic line, underscoring the detrimental consequences of co-pathology.^38^ Collectively, these in vivo findings demonstrate that Aβ and α-syn can engage in synergistic cross-interactions that influence aggregate formation, propagation, and downstream neuropathology. However, the molecular mechanisms underlying these heterotypic interactions and how they translate into the neuropathological features of mixed dementias remain largely undefined.

One potential consequence of these heterotypic interactions is the generation of distinct amyloid fibril conformations, or polymorphs. Amyloid fibril polymorphism can arise when aggregation occurs under different molecular environments or in the presence of cofactors such as pH, salts, lipids, nucleic acids, or other proteins.^40–49^ Because fibril conformation strongly influences biological activity, this structural diversity is thought to contribute to the heterogeneous neuropathological phenotypes observed across neurodegenerative diseases.^14,15,50–54^ Supporting this idea, morphologically distinct α-syn inclusions have been observed in singly versus doubly transgenic mouse brains,^38^ raising the possibility that heterotypic molecular interactions with Aβ may reshape α-syn fibril structure and thereby alter its pathological properties. Nevertheless, how Aβ specifically alters α-syn fibril conformation, and whether these structural changes translate into distinct neuronal pathologies, remains poorly understood.

To address this knowledge gap, we investigated how two distinct Aβ42 species, monomers and PFFs, differentially modulate α-syn aggregation kinetics and resulting fibril structure, and whether the Aβ42-modulated α-syn fibrils exhibit distinct pathological activities in neuronal cells. Our findings show that the aggregation state of Aβ42 is an important determinant of α-syn aggregation behavior: Aβ42 monomers and PFFs distinctly shape α-syn fibril formation, giving rise to structurally distinct α-syn fibril populations with markedly different capacities to induce intraneuronal α-syn aggregation. These results provide molecular insight into how heterotypic interactions between amyloidogenic proteins generate fibril structural diversity and contribute to the neuropathological complexity of mixed dementias.

## Materials and Methods

### α-synuclein (α-syn) Protein Expression and Purification

Full-length α-syn was expressed in *Escherichia coli (E. coli)* BL21 (DE3) competent cells. A single colony was inoculated into 10 mL of Luria-Bertani (LB) medium containing ampicillin (100 µg/mL) and incubated overnight at 37 °C with shaking at 150 rpm. The next day, the 10 mL culture was transferred into 1 L of autoclaved LB medium containing ampicillin (100 µg/mL) and incubated at 37 °C with shaking at 250 rpm until the optical density at 600 nm (OD_600_) reached 0.7-0.9. Protein expression was then induced by adding 1 mM isopropyl-β-D-1-thiogalactopyranoside (IPTG) and incubated at 37 °C with shaking at 250 rpm for 4 h. Cells were harvested by centrifugation at 7,000 rpm for 20 min and stored at -80 °C.

The cell pellet obtained from 1 L of LB culture was resuspended in 25 mL of lysis buffer (10 mM Tris-HCl, 1 mM EDTA, 1 mM PMSF, and half of Sigma Fast protease inhibitor cocktail tablet, pH 7.8). After transferring the resuspended cells into a centrifuge tube, lysozyme was added to a final concentration of 0.2 mg/mL, and the mixture was incubated on ice for 20-30 min.

Cells were lysed by sonication using a Branson Digital Sonifier SFX 250 at 40% power with a pulse cycle of 10 s on and 15 s off for a total sonication time of 3.5 min. The lysate was incubated on ice for 10-15 min, followed by centrifugation at 16,000 rpm for 1 h at 4 °C.

The milky supernatant was transferred to a transparent centrifuge tube for acid precipitation. pH was adjusted to 3.5 and stirred at room temperature for 20 min. The sample was centrifuged at 16,000 rpm for 30 min at 4 °C, resulting in a clear supernatant. The pH was then adjusted back to 7.5, and the sample was centrifuged at 16,000 rpm for 1 h at 4 °C to remove residual aggregates.

The collected supernatant was filtered through a PVDF syringe filter with 0.45 μm pore size (Avantor, VWR® Syringe Filters) and dialyzed overnight at 4 °C against 1 L of low-salt anionic exchange buffer A (10 mM Tris-HCl, 25 mM NaCl, 1 mM EDTA, pH 7.6) using a dialysis membrane with a molecular weight cutoff (MWCO) of 3.5 kDa (Spectrum Laboratories, Inc., Spectra/Por®3 Dialysis Membrane).

The following day, the dialyzed α-syn was filtered through a PES syringe filter with 0.22 μm pore size (Avantor, VWR® Syringe Filters) and loaded onto a double-stacked pre-packed HiTrap^TM^ Q Sepharose High Performance anion exchange chromatography column (Cytiva Sweden AB, #17115401) equilibrated with low-salt anionic exchange buffer A. α-Syn was eluted by gradually increasing the concentration of high-salt anionic exchange buffer B (10 mM Tris-HCl, 1 M NaCl, 1 mM EDTA, pH 7.6) from 0% to 100% over 100 mL. α-Syn-containing fractions were collected at a conductivity of approximately 30 mS/cm, concentrated to ∼14 mg/mL using a 10 kDa Amicon ®Ultra centrifugal filter device (Millipore), and dialyzed overnight at 4 °C against 1 L of size exclusion buffer (10 mM Tris-HCl, 100 mM NaCl, pH 7.6) using a dialysis membrane with a MWCO of 3.5 kDa (Spectrum Laboratories, Inc., Spectra/Por®3 Dialysis Membrane).

The dialyzed protein was loaded onto a pre-packed HiPrep 16/60 Sephacryl S-200 High Resolution preparative size exclusion chromatography column (Cytiva Sweden AB, #17116601) equilibrated with size exclusion buffer. α-Syn protein eluted between 40 mL and 60 mL of the total 126 mL elution volume. The purity of the collected fractions was assessed by SDS-PAGE followed by Coomassie blue staining (Blazin’ Blue^TM^ Protein Gel Stain, GOLD BIO, Laboratories). Protein concentrations were determined using a NanoDrop ND-1000 UV/Vis Spectrophotometer (Thermo Fisher Scientific). Purified α-syn was dialyzed overnight against 1 L of distilled water at 4 °C, concentrated to 1 mM, and stored at -80 °C.

All column chromatography was performed using an AKTA^TM^ pure (GE Healthcare Bio-Sciences AB, Sweden).

### Homotypic and Heterotypic Fibril Formation

Aβ42 (Beta-Amyloid (1-42), TFA, rPeptide) stock solution was prepared at 4mM in DMSO. Homotypic Aβ42 fibrils were generated by diluting the 4 mM Aβ42 stock to a final concentration of 40 µM in 10 mM Tris-HCl (pH 7.6) and incubating at 37 °C without agitation. Homotypic α-syn fibrils were prepared by diluting a 1 mM α-syn stock in DI water to a final concentration of 200 µM in 10 mM Tris-HCl (pH 7.6), followed by incubation at 37 °C without agitation.

Heterotypic fibrils were generated by incubating 200 µM α-syn monomer with 2, 4, or 8 mol % (relative to α-syn monomer) of Aβ42 monomers or Aβ42 preformed fibrils (PFFs) in 10 mM Tris-HCl (pH 7.6) at 37 °C without agitation. The resulting heterotypic fibrils formed with Aβ42 monomers are referred to as α-syn + 2M, α-syn + 4M, and α-syn + 8M, whereas those formed with Aβ42 PFFs are referred to as α-syn + 2P, α-syn + 4P, and α-syn + 8P.

All fibril formation was confirmed by negatively stained transmission electron microscopy (TEM).

### Preformed Fibrils (PFF) Formation

Preformed fibrils (PFF) were generated by sonicating the fibrils described above using a sonifier (Branson Digital Sonifier SFX 250 with Sonifier Sound Enclosure) at 10% amplitude power with a pulse cycle of 1 s on and 4 s off for a total sonication time of 2 min. The average PFF length (∼80 nm) was confirmed by negatively stained transmission electron microscopy. PFFs generated from the heterotypic fibrils α-syn + 2M, α-syn + 4M, α-syn + 8M, α-syn + 2P, α-syn + 4P, and α-syn + 8P are referred to 2M-PFF, 4M-PFF, 8M-PFF, 2P-PFF, 4P-PFF, and 8P-PFF respectively. All prepared PFFs were stored at -80 °C and thawed and bath-sonicated for 20 s immediately before use.

### Solid-State NMR (ssNMR) Sample Preparation

For ssNMR measurements, uniformly ^13^C, ^15^N-labeled α-syn protein was expressed in *E. coli* BL21 (DE3) competent cells. Cells were grown in 2 L of LB medium containing 100 µg/ml of ampicillin at 37 °C with shaking at 250 rpm until OD_600_ reached 0.5-0.6. Cells were harvested by centrifugation at 7,000 rpm for 20 min. The collected cells were resuspended in 1 L of autoclaved M9 medium containing 2 g of ^13^C-labeled glucose, 1 g of ^15^N-labeled ammonium chloride, 2 mM of MgSO_4_, 0.1 mM of CaCl_2_, 100 µg/ml of ampicillin, and 10 mL Gibco^TM^ 100x MEM vitamin solution (Thermo Fisher Scientific). The M9 medium containing the resuspended cells was further incubated until the OD₆₀₀ reached 0.8–0.9. Protein expression was then induced with 1 mM IPTG, and incubation was continued for 4 h under the same conditions. Cells were collected by centrifugation at 7000 rpm for 20 min and stored at -80 °C. Uniformly ^13^C-and ^15^N-labeled α-syn was purified using the same procedures as those used for unlabeled α-syn described above.

Heterotypic fibrils were generated by incubating uniformly ^13^C, ^15^N labeled α-syn monomers (200 µM) with 2 or 8 mol% unlabeled Aβ42 monomers or 2 mol% unlabeled Aβ42 PFFs in 10 mM Tris-HCl buffer (pH 7.6) and incubated at 37 °C without agitation. Fibril formation was confirmed by negatively stained TEM images.

Fibrils were pelleted by centrifugation at 50,000 rpm for 24 h at 4 °C using a Beckman SW 60Ti rotor. The pellets were slowly dried in a desiccator to achieve ∼50% water by mass and then packed into a 3.2 mm MAS rotor.

### Transmission Electron Microscopy (TEM)

Negatively stained fibrils or PFFs were prepared on lacey carbon-coated copper grids (Ted Pella, Inc., 01824). A 10 µL of diluted sample was deposited onto a grid and allowed to adsorb for 1-2 min. The excess solution was blotted off, washed with 10 µL of distilled water, blotted again, and stained with 10 µL of UranyLess EM stain solution (Electron Microscopy Sciences) for 10 s.

For immunogold electron microscopy, a 10 µL aliquot of fibrils was deposited onto lacey carbon-coated copper grids (Ted Pella, Inc., 01824) and allowed to absorb for 10 min. Excess sample was blotted, and this step was repeated once. Grids were incubated with 10 µL of blocking buffer (1% bovine serum albumin (BSA) in phosphate-buffered saline (PBS)) for 30 min at room temperature. Following blocking, grids were incubated with 15 µL of primary antibody (6E10 anti-Aβ, BioLegend, 803014; 1:50 in blocking buffer) for 2 h at room temperature. Grids were then washed three times with 10 µL blocking buffer for 2 min and incubated with 15 µL of secondary antibody (12 nm gold-conjugated goat anti-mouse IgG (H+L), Jackson Immuno Research; 1:20 in blocking solution) for 2 h at room temperature. Grids were subsequently washed three times with 10 µL blocking buffer for 2 min, followed by three times wash with 10 µL distilled water for 2 min. Grids were stained with 10 µL UranyLess EM stain (Electron Microscopy Sciences) for 20 s. TEM images were acquired using a JEOL 2100F field-emission transmission electron microscope at 120 kV with a Gatan Ultrascan CCD camera and DigitalMicrograph (GMS3) software (Gatan Inc.).

### Thioflavin T (ThT) Fluorescence Monitoring of Fibril Formation Kinetics

ThT (25 mM) was added to a freshly prepared solution of monomeric α-syn (200 µM) in the presence of 2, 4, 8 mol% (relative to α-syn monomer) of Aβ42 monomer or Aβ42 PFF in 10 mM Tris-HCl buffer (pH 7.6). 200 µM α-syn monomer, 2, 4, 8 mol% (relative to α-syn monomer) of Aβ42 monomer or Aβ42 PFF were included as controls. Measurements were performed in Greiner Bio-One 96-well microplates (655900). To prevent sample dehydration during measurements, 50 µL of 10 mM Tris-HCl was added to the outermost wells.

Fluorescence was measured using a Synergy™ HT Multi-Mode Microplate Reader (Gen5^TM^ & Gen5 Secure, BioTek Instruments) at 37 °C with excitation at 440 nm and emission at 485 nm. Measurements were taken at 30 min intervals with a gain setting of 35. Before each measurement, the plate was orbitally shaken for 1 min at a medium speed. Each condition was measured in triplicate. ThT fluorescence curves were quantitatively analyzed by fitting each ThT kinetic curve to a sigmoidal growth equation,^55–57^

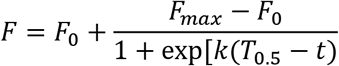

where F_0_ is the initial baseline, F_max_ is the final maximum fluorescence intensity after the growth phase has ended, k is the apparent fibril growth rate constant, and T_0.5_ is the time required to reach 50% of maximum ThT intensity. The T_lag_ was calculated as T_0.5_ − 2/k.

### Solid-State NMR (ssNMR) Experiments

All ssNMR experiments were conducted on an 800 MHz (18.8 Tesla) Bruker Advance III-HD NMR spectrometer equipped with a 3.2-mm ^1^H-^13^C-^15^N magic angle spinning (MAS) probe (Black Fox, LLC). ^13^C chemical shifts are reported on the TMS scale using the 38.48 ppm of CH_2_ signal of adamantane as a reference. All spectra were measured under 13.5 kHz MAS. 1D ^13^C direct polarization (DP) and 1D ^13^C cross polarization (CP) spectra were measured at 293K. 2D ^15^N-^13^C and ^13^C-^13^C correlation spectra were measured at 273 K. 2D ^13^C-^13^C dipolar-assisted rotational resonance (DARR) spectra were acquired with mixing times of 50 ms.^58,59^

#### Proteinase K Digestion

Proteinase K (PK) digestion was performed on heterotypic fibril mixtures, with homotypic α-syn fibrils used as a control. Fibrils were treated with PK to a final concentration of 0.05 mg/mL. After briefly vortexing, the samples were incubated at 37 °C without agitation for 10 min. The PK reaction was terminated by heating the samples at 90 °C for 15 min. Samples were lyophilized and resuspended in 200 µL of hexafluoroisopropanol (HFIP). The suspension was sonicated for 5 min in a water bath, vortexed for 30 s, and left capped in a fume hood overnight to ensure complete solubilization. The following day, HFIP was evaporated under a gentle stream of N_2_ gas. The dried proteins were resuspended in 50 µL denaturing buffer (8 M urea and 4% SDS), vortexed vigorously, sonicated for 5 min, and incubated at room temperature for 15 min with intermittent vortexing.

Protein digestion profiles were analyzed by SDS-PAGE, followed by silver staining using a commercial kit (Pierce^TM^ Silver Stain Kit, Thermo Fisher Scientific) according to the manufacturer’s instructions.

#### Cell Culture

To establish a dopaminergic neuron model, the human neuroblastoma cell line (SH-SY5Y; ATCC, CRL-2266) was selected and expanded in 25 cm^2^ flasks following the manufacturer’s protocols. To culture SH-SY5Y cells, Dulbecco’s Modified Eagle Medium (DMEM; 10-017-CV, Corning) was used as the basal growth medium, supplemented with 10% (v/v) heat-inactivated fetal bovine serum (hiFBS; SH30396.03, Cytiva), 1% (v/v) GlutaMAX-I (35050-06, Thermo Fisher) and 1% (v/v) penicillin-streptomycin (SV30010, Cytiva). SH-SY5Y cells at passages 3 to 10 were used as recommended by the cell supplier.

The process of dopaminergic neuronal differentiation started with trypsinization and resuspension of SH-SY5Y cells in growth medium. The harvested cells were then introduced into 48-well plates at a density of 2.5 × 10^5^ cells/mL. The initial culture was maintained in their growth medium to promote cell adherence and stabilization. 2 days after seeding, the SH-SY5Y cell growth medium was replaced with the first differentiation medium consisting of DMEM supplemented with 2.5% (v/v) hiFBS, 1% (v/v) GlutaMAX-I, 1% (v/v) penicillin-streptomycin, and 10 μM retinoic acid (50-165-6969, Fisher Scientific) to induce differentiation into dopaminergic neuronal cells. After 5 days, the first differentiation medium was switched to a second differentiation medium based on Neurobasal-A (10888022, Thermo Fisher) mixed with 50 ng/mL brain-derived neurotrophic factor (BDNF; P23560, FUJIFILM Irvine Scientific), 20 mM potassium chloride (KCl; 7447-40-7, Avantor Science), 1% (v/v) B27 (17-504-044, Fisher Scientific), 1% (v/v) GlutaMAX-I, and 1% (v/v) penicillin-streptomycin. The second differentiation medium was also maintained for an additional 5 days. Both the first and second differentiation media were refreshed every other day to support and sustain the dopaminergic differentiation of SH-SY5Y cells.

#### Treatment of α-syn PFFs

To investigate the pathological impacts of polymorphic α-syn fibrils, our dopaminergic neuronal cells were exposed to different α-syn PFF conformers. For this work, a 50 µM stock solution of each α-syn PFF conformer was sonicated for 15 s to promote uniform dispersion and achieve an optimal working size (∼100 nm). To treat dopaminergic neuronal cells, the sonicated stock solution was diluted to a final concentration of 1 μM in the second differentiation medium. On Day 12 of culture, this α-syn PFF-containing medium was added to each well containing dopaminergic neuronal cells. The cultures were then incubated with the fibrils for 48 h. After incubation, the medium was replaced with fresh differentiation medium. The treated cells were maintained for an additional 3 days before analysis.

#### Immunostaining

For immunostaining, dopaminergic neuronal cells were fixed with 4% paraformaldehyde (AR1068, Boster) for 30 min at room temperature (RT) and washed twice using Dulbecco’s Phosphate-Buffered Saline (DPBS). After fixation, the fixed cells were permeabilized with 0.1% Triton X-100 (9002-93-1, VWR) for 15 min. To prevent non-specific binding, the cells were blocked with 1.5% bovine serum albumin (BSA; 22013, Biotium) for 30 min at RT. Subsequently, the cells were incubated overnight at 4 °C with primary antibodies that selectively bind to the proteins of interest.

To visualize the actin cytoskeleton, Phalloidin-CF 430 (Biotium) was applied at a dilution of 1:100. Additionally, neuronal differentiation was demonstrated using primary antibodies of rabbit polyclonal anti-MAP2 (A16829, 1:200, Antibodies.com) for dendritic extensions, and mouse monoclonal anti-GAP43 (A85392, 1:5000, Antibodies.com) for axonal projections. To assess the dopaminergic characteristics of neuronal cells, Tyrosine Hydroxylase (TH) expression was examined using a mouse monoclonal anti-TH antibody (A104316, 1:500, Antibodies.com). For observing intracellular aggregation burdens, we used a rabbit monoclonal anti-α-synuclein aggregate [MJFR-14-6-4-2] (AB209538, 1:5000, Abcam) antibody.

After incubation with primary antibodies, the cells were washed twice with DPBS and incubated for 2-3 h at RT with fluorescently labeled secondary antibodies (A32732, ThermoFisher; A32723, ThermoFisher; A32731, ThermoFisher; A11032, ThermoFisher). Hoechst (33342, ThermoFisher) was also used for nuclear staining.

Finally, fluorescence images of the cells were captured using the Nikon Ti2-Eclipse microscope. α-syn fibrils labeled with MJFR-14-6-4-2 were imaged with excitation at 500 nm and emission at 535 nm. Exposure times were 100 ms for TH, and 200 ms for MJFR-14-6-4-2. Image processing was performed using NIS Elements software.

#### Quantification of intraneuronal α-syn aggregation

To quantitatively assess the data, the number of puncta-positive cells (nuclei) and total number of cells (nuclei) were counted and the punctated cell ratio (%) was calculated using the following equation:

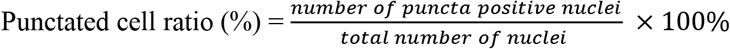

Puncta positive cell ratio was averaged from three samples for each experimental condition.

For quantification of mean punctate aggregation length, ten punctate structures were selected per sample, consisting of the three largest, the three smallest, and four of intermediate size. The major axis length was measured for each structure, given the typically elongated morphology of α-syn puncta. These ten measurements were averaged to obtain a representative value per sample, and the mean of three independent samples per condition was used as the final value. Punctate structures with dimensions close to or exceeding that of cell nuclei were excluded from analysis

### Statistical analysis

Statistical value of the obtained data was assessed by a two-tailed t-test. Data from three independent experiments were presented as mean ± SD.

## Results

### α-syn fibril formation in the presence of Aβ42 monomers and preformed fibrils (PFFs)

To examine how the presence of Aβ42 influences α-synuclein (α-syn) fibril formation, we monitored the aggregation kinetics of 200 µM α-syn using a thioflavin T (ThT) fluorescence assay in the absence or presence of Aβ42 monomers or Aβ42 preformed fibrils (PFFs) at 2, 4, or 8 mol% relative to α-syn, corresponding to Aβ42 concentrations of 4, 8, and 16 µM, respectively. The resulting heterotypic assemblies formed with Aβ42 monomers are referred to as α-syn+2M, α-syn+4M, and α-syn+8M, and those formed with Aβ42 PFFs as α-syn+2P, α-syn+4P, and α-syn+8P. As controls, Aβ42 monomers and PFFs at 4, 8, and 16 µM were analyzed independently to account for their contributions to the ThT fluorescence signal. Aβ42 monomers exhibited concentration-dependent increases in ThT fluorescence, beginning at approximately 12, 8, and 4 h for the 4, 8, and 16 µM samples, respectively, consistent with β-sheet-rich fibril formation (**Fig. S1A**). In contrast, Aβ42 PFF controls displayed elevated initial ThT intensities relative to the monomer controls, consistent with the presence of pre-existing β-sheet-rich fibrillar structures (**Fig. S1B**). Importantly, the ThT signals from all Aβ42 controls were substantially lower than those observed in α-syn-containing samples (overall less than 10%), confirming that the Aβ42 contribution to the overall ThT readout in co-aggregation experiments is negligible.

To quantitatively assess the concentration- and species-dependent effects of Aβ42 on α-syn aggregation, we extracted two kinetic parameters from the ThT curves, the lag time (T_lag_) and the half-time of aggregation (T_50_), by fitting the ThT fluorescence curves to a sigmoidal growth equation.^55–57^ These parameters were used as quantitative metrics for comparing aggregation kinetics across different conditions.

Analysis of the ThT aggregation profiles revealed that Aβ42 monomers and Aβ42 PFFs exert opposing effects on α-syn fibril formation (**Fig. 1**). Aβ42 PFFs exhibited a trend toward accelerated α-syn aggregation with increasing Aβ42 concentration, as reflected by gradual reductions in both T_lag_ and T_0.5_ (**Fig. 1D**). This behavior indicates that Aβ42 PFFs serve as heterologous nucleation templates for α-syn monomers, facilitating α-syn fibril assembly despite the absence of sequence homology between the two proteins. In contrast, Aβ42 monomers showed a concentration-dependent trend toward delaying α-syn fibril formation, with both T_lag_ and T_0.5_ trending to increase as the Aβ42 monomer concentration increased (**Fig. 1D**). This inhibitory effect suggests that Aβ42 monomers interfere with the homotypic intermolecular interactions among α-syn molecules that drive nucleation and subsequent fibril growth.

**Figure 1.**
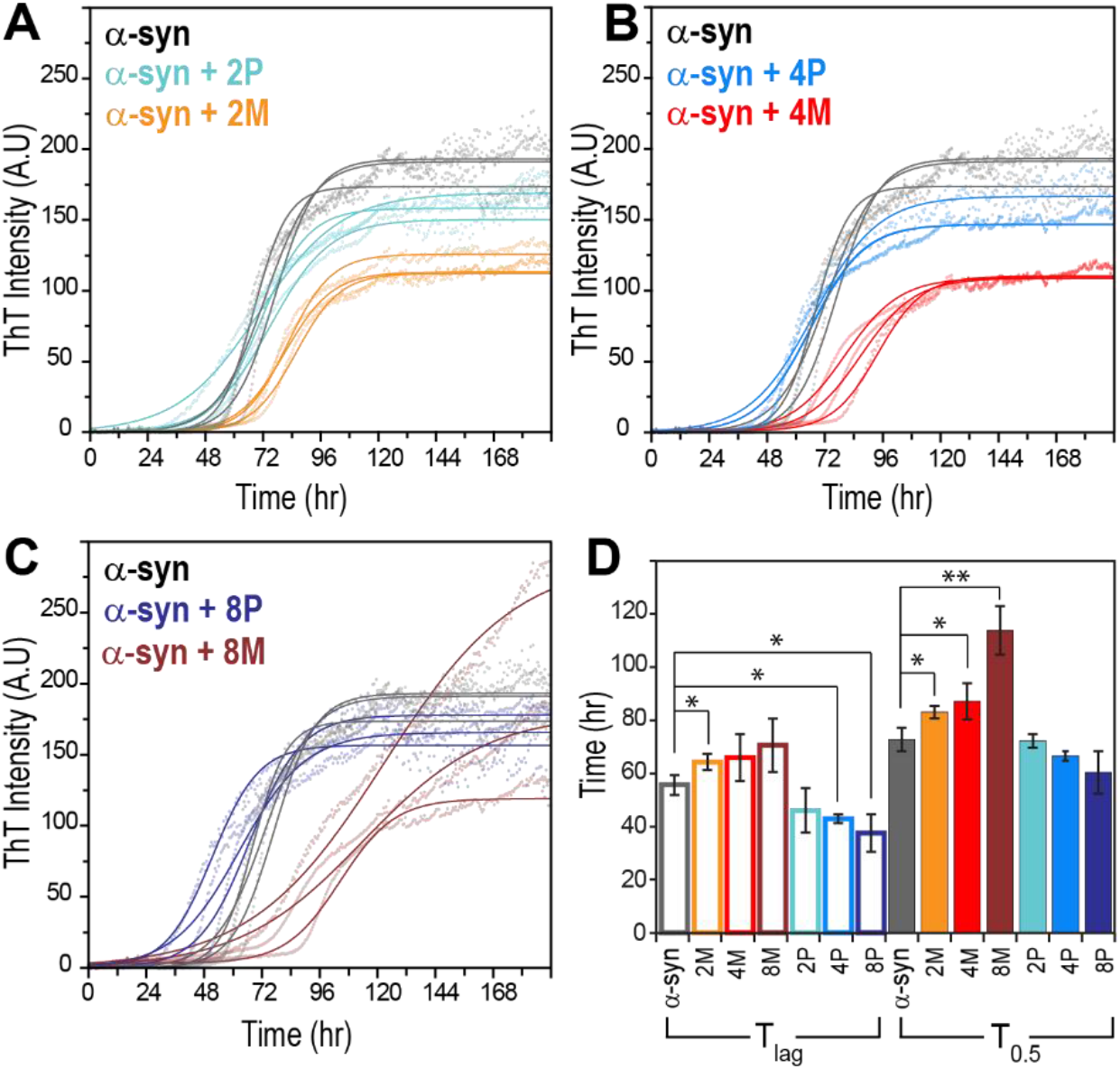
Concentration-dependent effects of Aβ42 PFFs and Aβ42 monomers on α-syn aggregation kinetics. (**A**) ThT fluorescence curves of α-syn aggregation kinetics for 200 µM α-syn monomer in the absence of Aβ42 (black) or in the presence of Aβ42 PFFs (P) and Aβ42 monomers (M) at 2 mol% (**A**; α-syn+2P, cyan; α-syn+2M, orange), 4 mol% (**B**; α-syn+4P, blue; α-syn+4M, red), and 8 mol% (**C**; α-syn+8P, dark blue; α-syn+8M, brown). Solid lines represent ThT curves fitted using the sigmoidal equation described in the Materials and Methods, whereas dotted lines represent the raw data. (**D**) Quantitative comparison of aggregation kinetics using T_lag_ (open bars) and T₀.₅ (solid bars), derived from the fitting of individual ThT curves using the sigmoidal equation in the Materials and Methods section. The same color scheme as in (**A-C**) is applied. Data show mean ± SD with n = 3. ***P < 0.001, **P < 0.01 *P < 0.05.

Then, to examine whether Aβ42 associates with α-syn fibrils during co-aggregation. we performed immunogold transmission electron microscopy (TEM) on α-syn fibrils grown in the presence of Aβ42 monomers and PFFs at different concentrations. The presence of Aβ42 within fibrils was probed using an antibody conjugated to 12 nm gold nanoparticles. While the pure α-syn fibrils showed the absence of gold nanoparticles near fibrils, the pure homotypic Aβ42 fibrils were shown to be decorated by gold particles, confirming the specificity to Aβ42 proteins of our labeling protocol using gold nanoparticle-conjugated antibodies (**Fig. S2**). α-Syn fibrils formed in the presence of either Aβ42 monomers or PFFs exhibited gold nanoparticle labeling along the fibrils (**Fig. S3**), indicating the presence of Aβ42 molecules associated with the fibrils. Although immunogold TEM cannot distinguish Aβ42 incorporation into the fibril core from Aβ42 bound to the fibril surface or located in close proximity, the observed labeling demonstrates that Aβ42 is closely associated with α-syn fibrils, supporting direct or near-direct interactions between the two proteins.

#### Conformationally different α-syn fibrils with Aβ42 monomers and PFFs

To examine whether Aβ42 monomers or PFFs alter α-syn fibril morphology, we acquired negative-stain TEM images of fibrils formed under each condition. α-Syn fibrils formed in the absence of Aβ42 predominantly exhibited straight, ribbon-like morphologies with no obvious periodic twisting (**Fig. 2A**).^60^ In contrast, α-syn fibrils formed in the presence of 2 or 8 mol% of Aβ42 monomers (α-syn+2M and α-syn+8M) contained substantial populations of twisted fibrils (**Figs. 2B** and **D**, yellow arrows), indicating that Aβ42 monomers can promote the formation of morphologically distinct α-syn fibrils. The crossover distance distributions of the twisted fibrils were centered at approximately 71.7 nm for α-syn+2M and 70.1 nm for α-syn+8M (**Figs. S4A** and **C**). Although the centers of these distributions were similar, α-syn+8M showed a somewhat broader distribution and a lower apparent abundance of twisted fibrils. In contrast, α-syn+4M displayed no obvious twisted fibrils (**Fig. 2C**, orange arrows), but displayed other morphologies, including fibrils composed of two apparent protofibrils with irregular twisting (magenta arrows). Thus, the morphological changes induced by Aβ42 monomers did not follow a simple concentration-dependent relationship.

**Figure 2.**
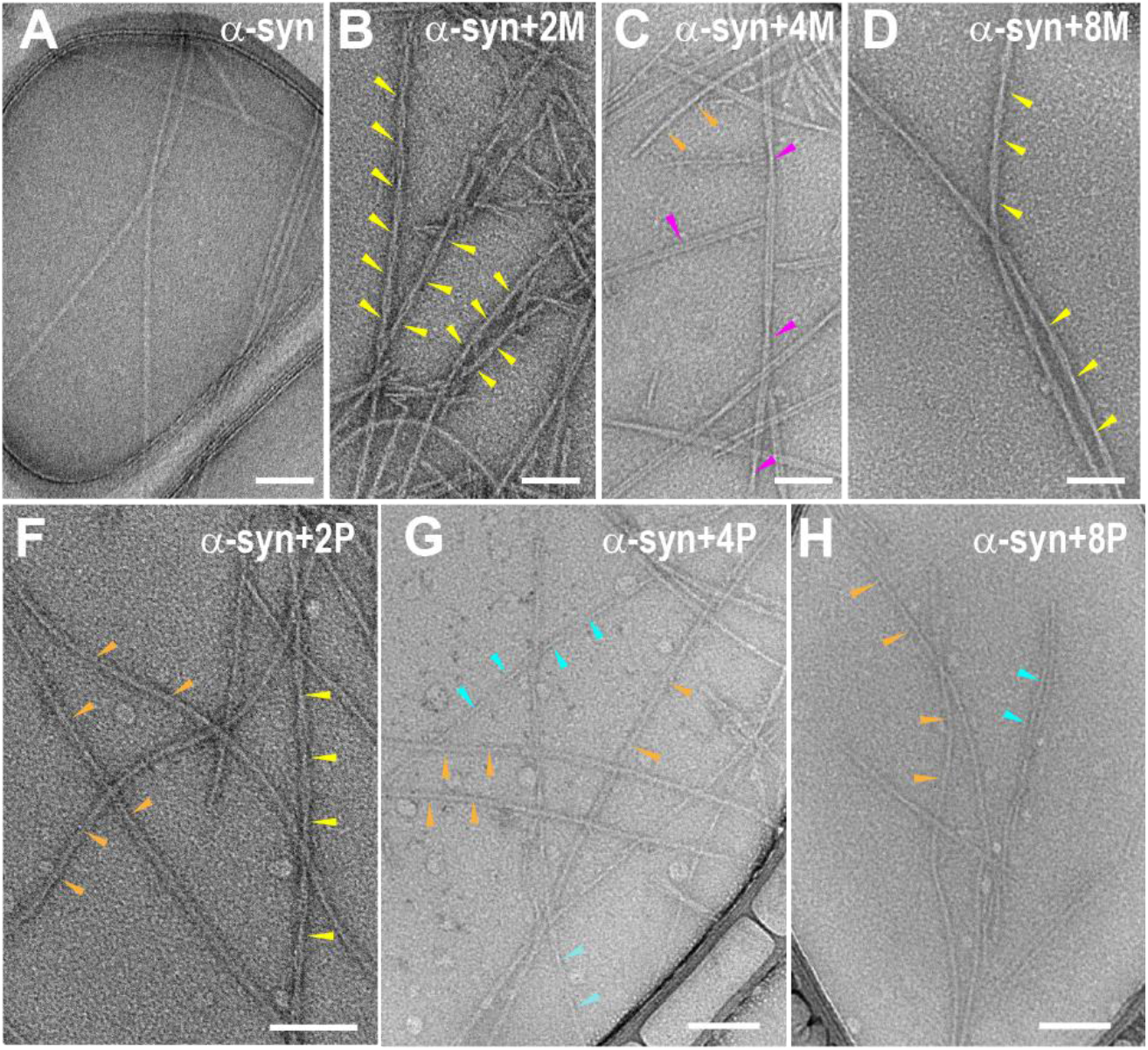
Concentration-dependent morphological changes in α-syn fibrils induced by Aβ42 monomers or PFFs. (**A**) Homotypic α-syn fibrils. (**B-D**) Heterotypic α-syn fibrils formed in the presence of 2, 4, or 8 mol% Aβ42 monomers, respectively. (**F-H**) Heterotypic α-syn fibrils formed in the presence of 2, 4, or 8 mol% Aβ42 PFFs, respectively. Yellow, magenta, orange, and cyan arrows indicate twisted fibrils, double-stranded fibrils, untwisted fibrils, and twisted narrower fibrils, respectively. Scale bars: 100 nm

Fibrils formed in the presence of Aβ42 PFFs also exhibited morphologies distinct from α-syn fibrils formed alone. Small populations of twisted fibrils were observed in the α-syn+2P, α-syn+4P, and α-syn+8P samples, although these populations appeared less abundant than those observed in α-syn+2M and α-syn+8M (**Figs. 2F-H**). The few twisted fibrils observed in α-syn+4P and α-syn+8P samples (cyan arrows) exhibited narrower fibril widths than the twisted fibrils observed in α-syn+2P or in the fibrils formed in the presence of Aβ42 monomers. Collectively, the TEM analysis demonstrates that both the aggregation state and concentration of Aβ42 present during α-syn fibril formation influence the resulting fibril morphology in a condition-dependent, rather than simply concentration-dependent manner. The coexistence of multiple morphologies within individual samples further suggests that Aβ42 can modulate the conformational heterogeneity of α-syn fibril populations.

Protein aggregates typically consist of a protease-resistant fibrillar core surrounded by a protease-susceptible disordered region. Structural differences among fibril polymorphs can therefore be assessed by comparing their proteinase K (PK) digestion patterns.^61–64^ To assess whether Aβ42 alters the protease-resistant architecture of α-syn fibrils, fibrils formed in the absence or presence of Aβ42 monomers or PFFs at different concentrations were subjected to PK digestion, and the resulting fragments were analyzed by SDS-PAGE (**Fig. S5**). PK digestion of α-syn fibrils formed in the absence of Aβ42 yielded two discrete low-molecular-weight fragments below 11 kDa together with a residual intact α-syn monomer band at ∼17 kDa. Under the same PK digestion condition, fibrils formed in the presence of Aβ42 exhibited digestion profiles that differed from the α-syn-only sample and from one another. Among the Aβ42 PFF-containing samples, α-syn+2P and α-syn+8P fibrils showed little or no detectable full-length α-syn band and displayed distinct distributions of lower-molecular-weight PK-resistant fragments, whereas α-syn+4P retained a band near ∼17 kDa together with additional lower-molecular-weight fragments. Fibrils formed in the presence of Aβ42 monomers also exhibited condition-specific digestion patterns, with α-syn+2M showing a profile distinct from those of α-syn+4M and α-syn+8M. Although the PK digestion patterns showed no clear dependence on Aβ42 concentration or aggregation state, the observed differences in protease susceptibility suggest that Aβ42 modulates α-syn fibril conformation in a condition-specific manner rather than through a simple concentration-or species-dependent effect.

To examine Aβ42-induced structural changes in α-syn fibrils at the molecular level, we measured solid-state NMR (ssNMR) spectra of uniformly ^13^C, ^15^N-labeled α-syn fibrils formed in the absence of Aβ42 or in the presence of 2 mol% Aβ42 monomers (α-syn+2M), 2 mol% Aβ42 PFFs (α-syn+2P), or 8 mol% Aβ42 monomers (α-syn+8M). One-dimensional ^13^C spectra were acquired using direct-polarization (DP) or cross-polarization (CP), which preferentially detect signals from dynamically disordered and conformationally rigid regions within fibrils, respectively.^12,65–67^ All four samples exhibited relatively strong signals in both DP and CP spectra at 273K, indicating that all fibrils contain both a rigid fibril core and a dynamic disordered region (**Fig. S6**). Importantly, each sample displayed a distinguishable spectral profile in both CP and DP spectra, with clear differences in peak positions and relative intensities, particularly in the Cα and carbonyl (CO) regions (highlighted by red boxes in **Fig. S6**). These spectral differences indicate that each aggregation condition generates a structurally distinct α-syn fibril population. Notably, α-syn+2M fibrils exhibited the sharper and better-resolved resonances in the CP spectrum than the other samples, suggesting a more conformationally homogeneous rigid fibril population.

Two-dimensional ^13^C-^13^C and ^15^N-^13^C spectra provide further evidence for Aβ42-dependent conformational differences in α-syn fibrils (**Fig. 3**). Relative to homotypic α-syn fibrils, α-syn+2P, α-syn+2M, and α-syn+8M all exhibited significant chemical shift changes, including prominent differences involving Gly, Ala, and Thr residues (**Figs. 3A-D**). Notably, the α-syn fibrils formed in the presence of Aβ42 monomers displayed more extensive spectral changes in the 2D ^15^N-^13^C spectra than those formed with Aβ42 PFFs. Together, the 1D and 2D ssNMR data demonstrate that the molecular conformation of α-syn fibrils is sensitive to both the aggregation state of Aβ42 and its concentration, resulting in distinct α-syn fibril conformational ensembles.

**Figure 3.**
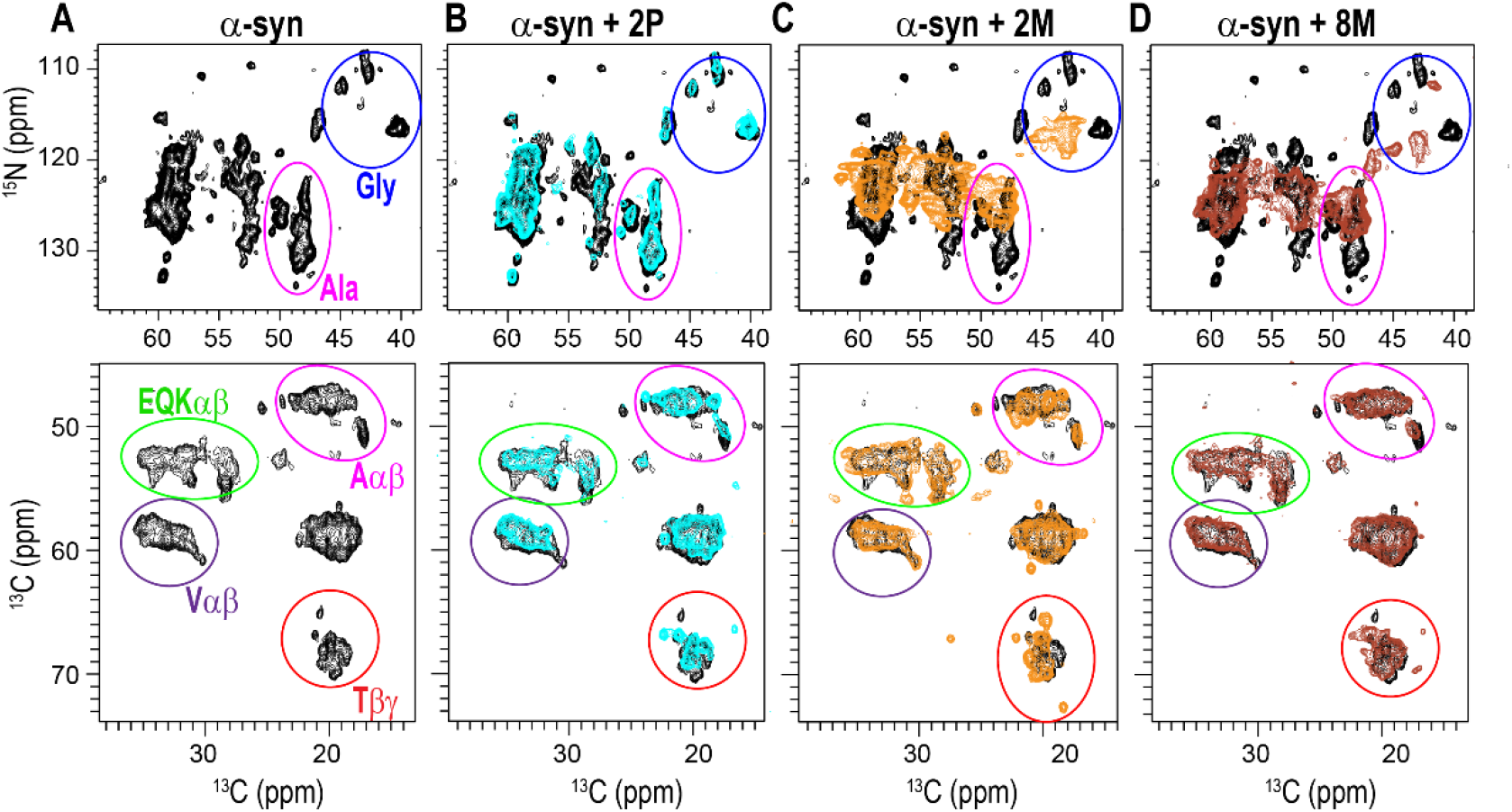
Aβ42-induced conformational changes in α-syn fibrils revealed by ssNMR. 2D ^15^N-^13^C (top) and ^13^C-^13^C (bottom) spectra of α-syn fibrils formed without (**A**, Black) or with 2 mol% Aβ42 PFFs (**B**, Cyan), 2 mol% Aβ42 monomers (**C**, Orange), or 8 mol% Aβ42 monomers (**D**, dark red). Heterotypic fibril spectra are overlaid with homotypic α-syn fibrils (Black). Colored circles indicate characteristic chemical shift regions for Thr (red), Ala (magenta), Gly (blue), Val (purple), and Glu/Gln/Lys (green) residues.

Taken together, the TEM analysis, PK digestion, and ssNMR spectra demonstrate that the presence of Aβ42 during α-syn fibril formation generates structurally distinct α-syn fibril populations. These differences are reflected in fibril morphology, protease susceptibility, and molecular-level ssNMR spectral patterns. Although the structural variations do not follow a simple relationship with Aβ42 concentration or aggregation state across all measurements, the combined data indicate that both monomeric and fibrillar Aβ42 can modulate α-syn fibril conformation in a condition-dependent manner.

#### Distinct intraneuronal aggregation induced by heterotypic α-syn fibrils

Given that α-syn fibril conformation is a key determinant of pathological outcomes in synucleinopathies,^68–70^ we next asked whether the polymorphic α-syn fibrils generated through heterotypic interactions with distinct Aβ42 species elicit divergent neuropathological effects. To test this, we generated six α-syn PFF variants by sonicating α-syn fibrils formed in the presence of 2, 4, or 8 mol% Aβ42 monomers or Aβ42 PFFs (2M-PFF, 4M-PFF, 8M-PFF, 2P-PFF, 4P-PFF, and 8P-PFF, respectively). TEM measurements of the six Aβ42-modulated PFF preparations and the homotypic α-syn PFF control confirmed comparable post-sonication length distributions, with mean seed lengths of 76.4-83.6 nm for the Aβ42-modulated samples and 81.9 nm for homotypic α-syn PFFs (**Table S1**). These Aβ42-modulated α-syn PFF variants were then applied to our neuronal cell model to examine how their distinct conformations shape α-syn pathology. For this work, we focused on Parkinson’s disease dementia (PDD) as our model disease, which is frequently associated with the co-occurrence of Aβ and α-syn aggregates and therefore provides a pathologically relevant context to study heterotypic interactions between these two proteins. Accordingly, we employed our cellular model of dopaminergic neurons, a cell type directly relevant to the motor deficits observed across PD and PDD.

To generate the midbrain neuronal population in vitro, we differentiated human neuroblastoma cells into dopaminergic neurons using our established protocol.^60,71^ Briefly, the cells were seeded into 48-well plates and cultured to confluency (**Fig. 4A**), and neuronal differentiation was then induced through sequential application of two distinct differentiation media formulations designed to drive morphological maturation (**Fig. 4B**). Early in differentiation, the cells began to extend thin projections (**Fig. 4B**, Day 2), which gradually developed into a dense network of neurite-like structures at later time points (**Fig. 4B**, Day 10). After 11 days of differentiation, the cells were evaluated for neuronal features by immunostaining for the dendritic marker microtubule-associated protein 2 (MAP2) and the axonal marker growth-associated protein 43 (GAP43). Robust expression of both neuronal markers confirmed that the differentiated neuroblastoma cells had acquired characteristic neuronal phenotypes (**Fig. 4C**). Their dopaminergic identity was also validated by immunofluorescent staining for tyrosine hydroxylase (TH), an enzyme essential for dopamine biosynthesis and a well-established marker of midbrain dopaminergic neurons (**Fig. 4C**).

**Figure 4.**
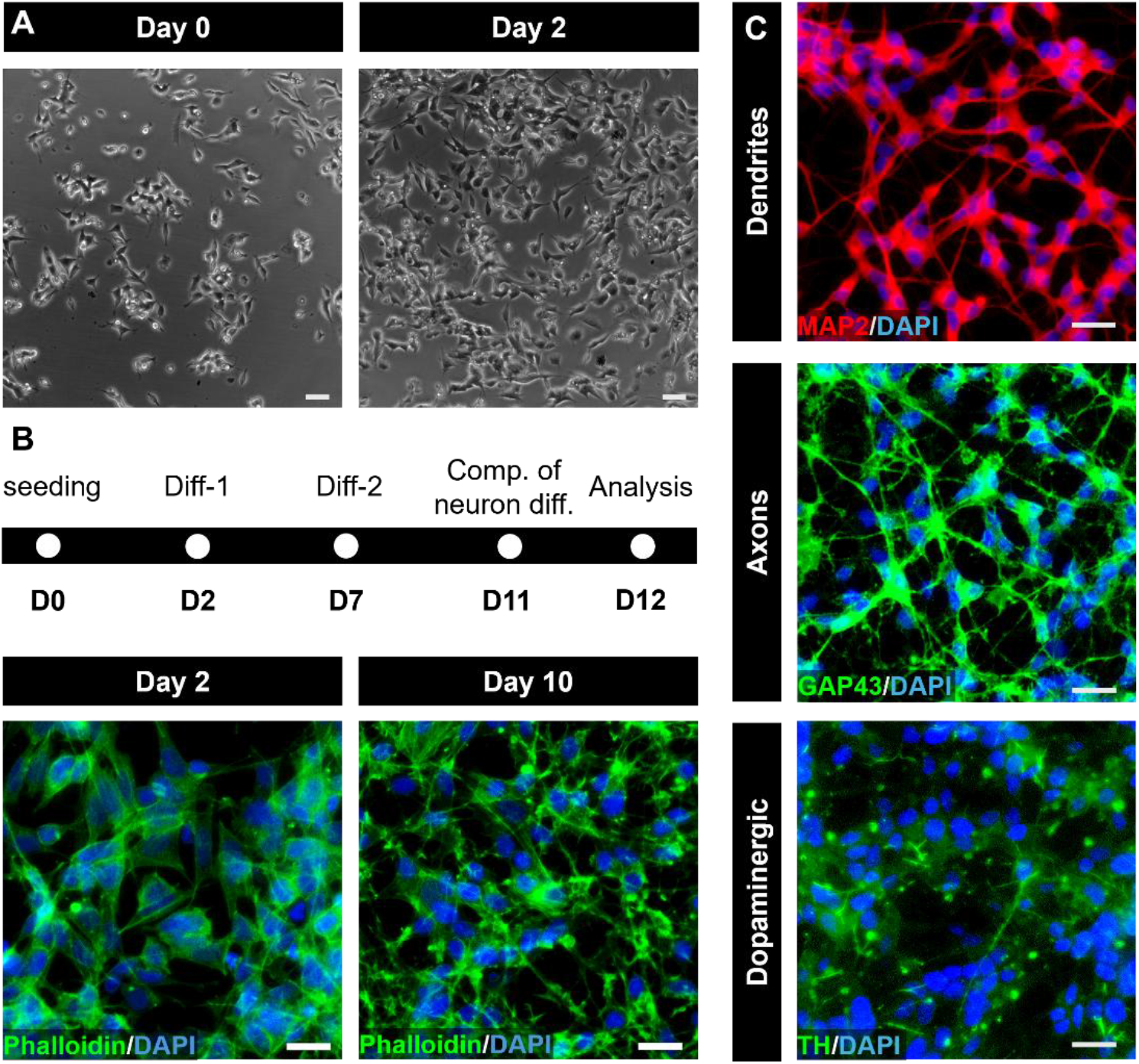
Establishment of a dopaminergic neuronal model. (**A**) Neuroblastoma cells prior to dopaminergic neuronal differentiation. Scale bars: 100 µm. (**B**) Timeline of dopaminergic differentiation for neuroblastoma cells. Abbreviations: D, day; comp., completion; diff., differentiation. Phalloidin staining shows morphological changes in neuroblastoma cells during differentiation at Days 2 and 10. Scale bars: 100 µm. (**C**) Immunofluorescence staining of differentiated neuroblastoma cells after 11 days confirms neuronal maturation and dopaminergic identity. MAP2 and GAP43 staining reveal dendritic and axonal structures, respectively, while tyrosine hydroxylase (TH) staining confirms the dopaminergic phenotype. Scale bars: 50 µm.

Using this neuronal platform, we then treated the dopaminergic neurons with each heterotypic α-syn PFF variant and assessed conformation-dependent effects on intraneuronal α-syn aggregation, a defining hallmark of the PD spectrum.^72,73^ As negative controls, neurons were either left untreated or treated with Aβ42 PFFs or Aβ42 monomers alone at the corresponding molar ratios (2, 4, or 8 mol% of α-syn), while α-syn PFFs generated in the absence of Aβ42 species served as a positive control. All homotypic and heterotypic α-syn PFFs were applied at 1 μM, a concentration previously shown to reliably template intraneuronal aggregation without inducing acute toxicity.^74–77^

Following 48 h of treatment, intraneuronal α-syn aggregation was assessed by immunofluorescent staining to determine whether the heterotypic α-syn PFF variants differed in their prion-like capacity to template de novo formation of α-syn aggregates in our dopaminergic neuronal cells. As shown in **Figs. 5** and **6**, PFF-treated groups exhibited discernible punctate structures indicative of misfolded α-syn aggregates, whereas negative control groups showed no detectable punctate aggregation. This overall increase with PFF treatment was further confirmed by quantification of the puncta-positive cell ratio and mean punctate aggregation length. Beyond this general effect, distinct α-syn PFF variants induced significantly different levels of aggregation in dopaminergic neuronal cells (**Figs. 5** and **6**). Among the variants generated through heterotypic interactions with Aβ42 PFFs, 8P-PFF induced the highest level of intraneuronal α-syn puncta, followed by 2P-PFF, whereas 4P-PFF produced the lowest aggregation burden. Notably, even the aggregation induced by 4P-PFF remained significantly elevated relative to both untreated controls and Aβ42 PFF-only controls across all corresponding molar ratios (**Fig. 5A**). Aggregation induced by homotypic α-syn PFF treatment was similar to that induced by 2P-PFF (**Fig. 5A**). These inter-variant differences were further supported by quantification of the puncta-positive cell ratio and mean punctate aggregation length (**Figs. 5B** and **C**).

**Figure 5.**
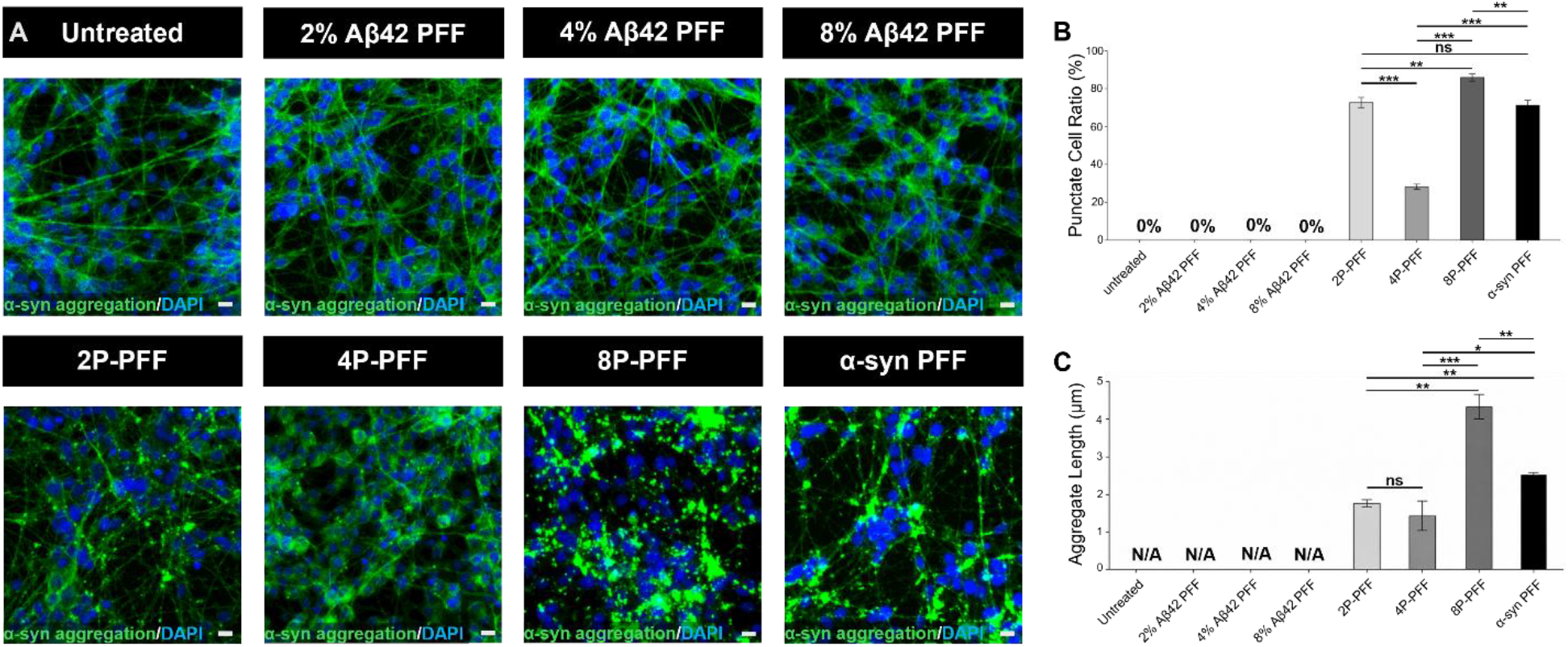
Intraneuronal α-syn aggregation induced by structurally distinct α-syn PFF conformers generated through heterotypic interactions with different concentrations of Aβ42 PFFs. (**A**) Immunofluorescence staining for MJFR-14-6-4-2 (green) was used to detect and visualize intracellular α-syn aggregates in dopaminergic neuronal cells under the indicated conditions. Scale bars: 10 µm. (**B**) Quantification of the puncta-positive cell ratio across heterotypic α-syn PFF variants and control groups. (**C**) Quantification of the mean punctate aggregation length across heterotypic and homotypic α-syn PFF-treated and control groups. Data are presented as mean ± SD. *P < 0.05, **P < 0.01 and ***P < 0.001 (n = 3).

**Figure 6.**
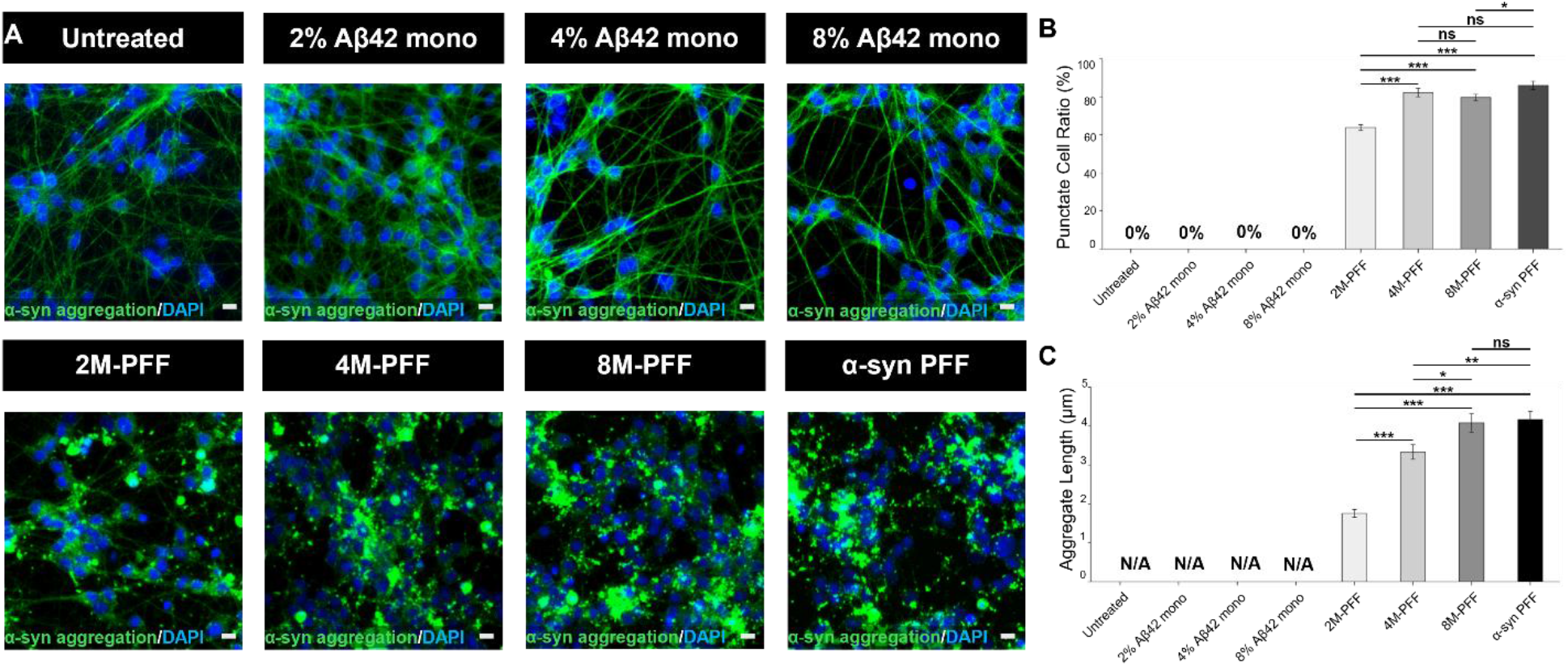
Different intraneuronal aggregation profiles induced by heterotypic α-syn PFF conformers generated through interactions with different concentrations of Aβ42 monomers. (**A**) α-syn aggregates in dopaminergic neuronal cells were visualized by immunofluorescence staining with MJFR-14-6-4-2 (green). Scale bars: 10 µm. (**B**) Quantification of the puncta-positive cell ratio across heterotypic α-syn PFF variants and controls. (**C**) Mean punctate aggregation length across heterotypic and homotypic α-syn PFF-treated and control groups. Data are presented as mean ± SD. *P < 0.05, **P < 0.01 and ***P < 0.001 (n = 3).

We then asked whether α-syn PFFs generated through heterotypic interactions with Aβ42 monomers also differed in their capacity to template intraneuronal aggregation in dopaminergic neurons. Among these variants, 8M-PFF, 4M-PFF, and homotypic α-syn PFFs produced similarly high levels of aggregation burden, whereas 2M-PFF induced the lowest (**Fig. 6A**). Quantification of both the puncta-positive cell ratio and mean punctate aggregation length confirmed this qualitative observation (**Figs. 6B** and **C**). Specifically, 8M-PFF, 4M-PFF, and homotypic α-syn PFFs showed largely comparable values across both metrics, except that 4M-PFF exhibited a shorter mean punctate aggregation length than 8M-PFF and homotypic α-syn PFFs (**Figs. 6B** and **C**). For both metrics, 2M-PFF exhibited the lowest values (**Figs. 6B** and **C**).

While the prior analysis clearly identified 2M-PFF as the variant with the lowest capacity to induce intraneuronal aggregation, the similar aggregation profiles of 8M-PFF, 4M-PFF, and homotypic α-syn PFF at 1.0 μM raised the question of whether these distinct conformers possess comparable aggregation-inducing capacities or whether the high dose saturated the cellular aggregation response, thereby masking underlying differences. To address this possibility, we repeated the experiment at a reduced concentration of 0.5 μM, reasoning that a sub-saturating dose may provide greater sensitivity for detecting inter-variant differences in intraneuronal aggregation-inducing capacity.

Unlike the comparable aggregation profiles observed at 1.0 μM, the reduced concentration revealed clear inter-variant differences in aggregation burden. Homotypic α-syn PFFs induced the highest aggregation burden, characterized by large, densely distributed punctate structures throughout the neuronal cells. 8M-PFF produced similarly widespread puncta, but of noticeably smaller size, whereas 4M-PFF generated puncta comparable in size to those induced by 8M-PFF but at a lower density. 2M-PFF consistently exhibited the lowest aggregation burden across all measures (**Fig. S7A**). Quantification of both the puncta-positive cell ratio and mean punctate aggregation length supported these qualitative observations (**Figs. S7B** and **C**). Both metrics confirmed the overall aggregation profile comparison, with homotypic α-syn PFFs showing the highest values, 4M-PFF and 8M-PFF exhibiting comparable intermediate levels, and 2M-PFF the lowest (**Figs. S7B** and **C**).

Collectively, these data from a human dopaminergic neuronal model demonstrate that α-syn PFF variants generated through heterotypic interactions with distinct Aβ42 species differ in their capacity to template intraneuronal α-syn aggregation. Notably, the two quantitative measures, puncta-positive cell ratio and mean punctate aggregation length, revealed distinct inter-variant patterns, indicating that fibril conformation differentially shapes multiple aspects of intraneuronal α-syn aggregation.

## Discussion

Although α-syn and Aβ are predominantly expressed and localized in distinct brain regions and subcellular compartments, accumulating evidence suggests that they can coexist within the same cellular environment. For example, Aβ has been shown to accumulate intracellularly through both endogenous production and neuronal uptake.^24,25^ Conversely, α-syn can be secreted into the extracellular space and subsequently taken up by neighboring cells, facilitating its propagation between neurons.^26^ These observations provide opportunities for direct heterotypic interactions between α-syn and Aβ during aggregation. Consistent with this possibility, numerous neuropathological studies have reported the coexistence of α-syn and Aβ aggregates in the same brain regions of patients with neurodegenerative diseases. For example, amyloid plaques and α-syn aggregates are observed together in the cortex of AD patients, and Lewy bodies (LBs) are found in the amygdala of AD brains.^20,21^ Moreover, the coexistence of these protein aggregates is associated with enhanced neuroinflammation, accelerated neurodegeneration, and increased pathological burden compared with either pathology alone.^32,34,35^ Together, these observations suggest that heterotypic interactions between α-syn and Aβ may play an important role in disease progression and underscore the need to understand how Aβ influences α-syn aggregation.

In our study, we found that Aβ42 modulates α-syn aggregation in an aggregation state-dependent manner: Aβ42 monomers delayed α-syn fibril formation, whereas Aβ42 PFFs promoted α-syn aggregation. Interestingly, reciprocal effects have also been reported for Aβ aggregation, where α-syn PFFs accelerate Aβ fibrillization, whereas α-syn monomers inhibit it.^78^ Together, these observations suggest that preformed fibrillar assemblies of one amyloid protein can catalyze the aggregation of another through heterotypic nucleation, indicating that fibril nucleation is not restricted to homotypic protein-protein interactions. Consistent with this model, a recent study demonstrated that early Aβ42 aggregates serve as nucleation sites for α-syn condensation, promoting the recruitment of α-syn molecules without altering the liquid-to-solid phase transition of α-syn.^79^ Our ThT kinetic analysis is in good agreement with this mechanism, as Aβ42 PFFs produced a more pronounced reduction in T_lag_ than in T_0.5_ (**Fig. 1D**), suggesting that fibrillar Aβ42 primarily facilitates the nucleation phase of α-syn aggregation rather than the subsequent fibril elongation process.

The delayed α-syn aggregation observed in the presence of Aβ42 monomers suggests that heterotypic interactions between soluble α-syn and Aβ42 may compete with the homotypic α-syn interactions required for nucleation and fibril growth. Consistent with this interpretation, co-oligomer formation between Aβ42 and α-syn has been reported to be more favorable than α-syn self-oligomerization.^80^ Previous studies have also identified interactions between the hydrophobic region of Aβ42 and the non-amyloid-β component (NAC) region of α-syn, both of which are critical for homotypic self-assembly of their respective proteins.^81^ Binding of Aβ42 monomers to α-syn NAC region could therefore sequester aggregation-competent α-syn monomers or mask intermolecular contact sites required for homotypic assembly. This model is consistent with our observation that the inhibitory effect became more pronounced as the concentration of Aβ42 monomers increased.

Although Aβ42 monomers and PFFs exerted opposite effects on α-syn aggregation kinetics, both forms of Aβ42 altered the structural properties of the resulting α-syn fibrils, as demonstrated by TEM, proteinase K digestion, and ssNMR analyses. These structural differences, in turn, appear to translate into distinct capacities of the Aβ42-modulated α-syn fibrils to induce intraneuronal α-syn aggregation in our dopaminergic neuronal model (**Figs. 5** and **6**), suggesting that fibril conformation may be an important determinant of downstream intracellular aggregation. However, the structural characterization in the present study does not resolve the three-dimensional molecular structure of each fibril polymorph. High-resolution structural studies, such as cryo-EM or detailed ssNMR-based structure determination, will therefore be required to define the specific conformational features responsible for the differences in intracellular aggregation. Nevertheless, our findings raise the possibility that heterotypic interactions between α-syn and Aβ42, occurring where the two proteins coexist in neurodegenerative disease, may generate conformationally distinct α-syn fibrils that differentially shape the trajectory of α-syn pathology and, by extension, downstream disease progression.

It should also be noted that aggregation-inducing capacity did not scale proportionally with Aβ42 concentration among either the PFF-derived or monomer-derived fibrils, further supporting the conclusion that fibril conformation, rather than Aβ42 dose, governs aggregation propagation capacity. Among the PFF-derived fibrils, 2P-PFF induced greater intracellular aggregation burden than 4P-PFF (**Fig. 5**). A similar lack of Aβ42 dose dependence was observed among the monomer-derived fibrils. Although 2M-PFF consistently produced the lowest aggregation burden among the three heterotypic variants, followed by 4M-PFF, none of the heterotypic α-syn PFF variants induced intracellular aggregation exceeding that produced by homotypic α-syn PFFs (**Figs. 6** and **S6**). Therefore, the absence of an Aβ42 concentration-proportional relationship across both fibril groups suggests that the amount of Aβ42 present during α-syn fibril formation is not the primary factor driving downstream aggregation-inducing activity. This interpretation is further supported by the finding that Aβ42 species alone, applied at ratios matching each heterotypic α-syn PFF preparation, produced no detectable intracellular α-syn aggregation (**Figs. 5** and **6**). This result excludes the possibility that residual Aβ42 itself, rather than the conformation it imparts to α-syn fibrils, accounts for the observed differences in intraneuronal aggregation. In sum, these results indicate that the conformational properties of α-syn PFFs, shaped by both the concentration and aggregation state of Aβ42 present during fibril formation, constitute a more critical determinant of intraneuronal aggregation-inducing activity than the amount of Aβ42 present alone.

By capturing conformation-dependent differences in aggregation burden, our quantitative analyses of puncta-positive cell ratio and mean aggregation length demonstrate that this dopaminergic neuronal model is sensitive enough to distinguish the cellular consequences of subtle structural variation among α-syn fibril polymorphs. This capacity makes the model a useful bridge between structural characterization and cellular pathology, offering a tractable readout for how fibril conformation translates into α-syn pathology at the cellular level. However, the current data cannot distinguish between two possible sources of this variation. Different conformations of fibrils may differ in how efficiently they are taken up by neurons, in how efficiently they template new aggregation once internalized, or in some combination of both. Resolving this question, and more broadly establishing a mechanistic link between fibril structure and prion-like seeding activity in a physiologically relevant context, will require further study. Even without that resolution, however, the present findings establish that the structure of heterotypic α-syn fibrils shapes their capacity to drive intracellular aggregation, a process central to synucleinopathies^5,50,82–84^ and relevant to mixed pathology in AD brains, where α-syn pathology frequently coexists with amyloid pathology.^85–87^

## Conclusion

Considering the heterogeneous molecular environments of the brain, where multiple amyloidogenic proteins coexist and can influence one another’s assembly, defining heterotypic protein interactions is essential for understanding neurodegenerative amyloid fibril formation. Our findings demonstrate that Aβ42 modulates α-syn aggregation in an aggregation state-dependent manner and alters the structural properties of the resulting α-syn fibrils. These Aβ42-modulated fibril polymorphs further exhibited distinct capacities to induce intraneuronal α-syn aggregation, suggesting that heterotypic interactions can influence not only fibril formation and conformation but also the capacity of the resulting fibrils to trigger intraneuronal aggregation. Together, these results highlight intermolecular interactions among amyloidogenic proteins as an important source of fibril structural and pathological heterogeneity and provide insight into how mixed protein pathologies may contribute to the accumulation, propagation, and overlap of pathological features across neurodegenerative diseases.

## Author Contributions

P.S^§^ and T.L^§^ contributed equally to this work.

J.P and M.L designed the project. P.S, T.L, and A.A performed experiments. M.L, J.P, P.S, and T.L wrote the manuscript.

## Notes

The authors declare no competing financial interest.

## Data Availability

2D ssNMR spectra and microscope images are available at 10.17632/djx4m8bggg.1.

Previously published ssNMR spectra of α-syn fibrils formed in the absence of Aβ42 were used for this work (published in Baek et al.^60^).

## Acknowledgements

We thank Drs. Ivan Hung and Zhehong Gan at the National High Magnetic Field Laboratory for their assistance with ssNMR experiments. The National High Magnetic Field Laboratory is supported by the National Science Foundation through NSF/DMR-2128556 and the state of Florida.

This work was supported by NIH grant NS139178 to M.L and J.P.

