## Supplemental Figures for "Aβ42-Driven α-synuclein Fibril Polymorphism and Distinct Intracellular Aggregation"

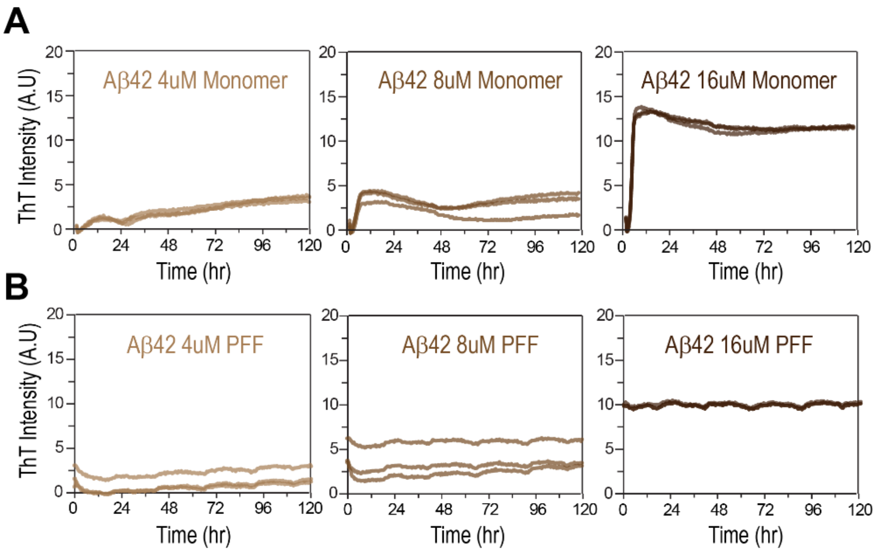


**Figure S1.** ThT controls. Concentration-dependent aggregation kinetics of Aβ42 monomers and pre-formed fibrils (PFFs) alone at 2, 4, and 8 mol% relative to 200 µM α-syn, corresponding to 4, 8, and 16 µM Aβ42. (**A**) Aβ42 monomer controls exhibited increased ThT fluorescence after approximately 12, 8, and 4 h at concentration of 4, 8 and 16 µM, respectively. (**B**) Aβ42 PFF controls show elevated initial ThT fluorescence, consistent with pre-existing β-sheet-rich fibrillar structures.

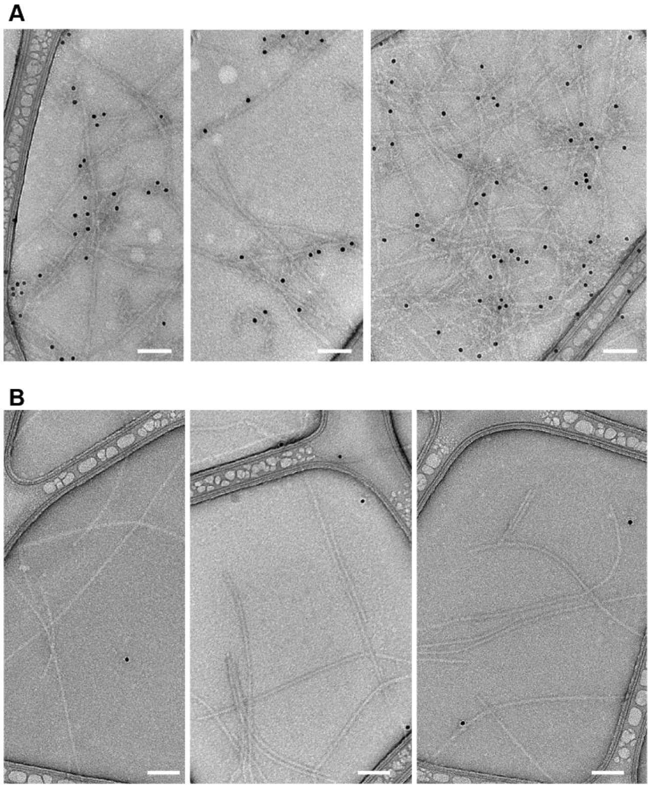


**Figure S2.** Immunogold TEM analysis of homotypic fibrils. Negative-stain transmission electron microscopy (TEM) of homotypic (**A**) Aβ42 fibrils and (**B**) α-syn fibrils labeled with an Aβ42-specific antibody conjugated to 12 nm gold nanoparticles (black dots). Gold particles are observed only on Aβ42 fibrils and not on α-syn fibrils, confirming the specificity of the antibody labeling. Scale bars: 100 nm.

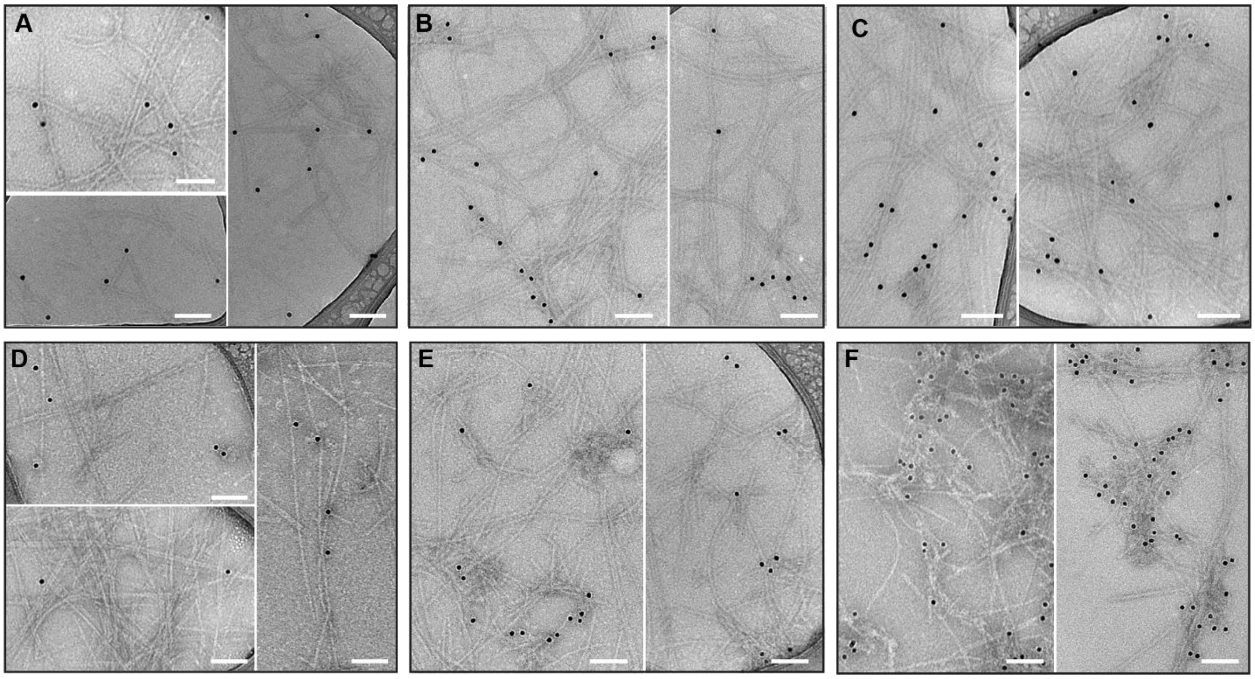


**Figure S3.** Immunogold TEM analysis of α-syn fibrils formed in the presence of Aβ42 monomers or PFFs. Negative-stain transmission electron microscopy (TEM) of heterotypic fibrils generated from 200 µM α-syn monomers with Aβ42 monomers at 2mol% (**A**; α-syn+2M), 4 mol% (**B**; α-syn+4M), and 8 mol% (**C**; α-syn+8M), or with Aβ42 PFFs at 2 mol% (**D**; α-syn+2P), 4 mol% (**E**; α-syn+4P), and 8 mol% (**F**; α-syn+8P). Black dots represent 12 nm gold nanoparticles conjugated to antibodies specific for Aβ42. Gold labeling associated with or adjacent to the fibrils indicates the presence of Aβ42 in close association with the fibrils. Scale bars: 100 nm.

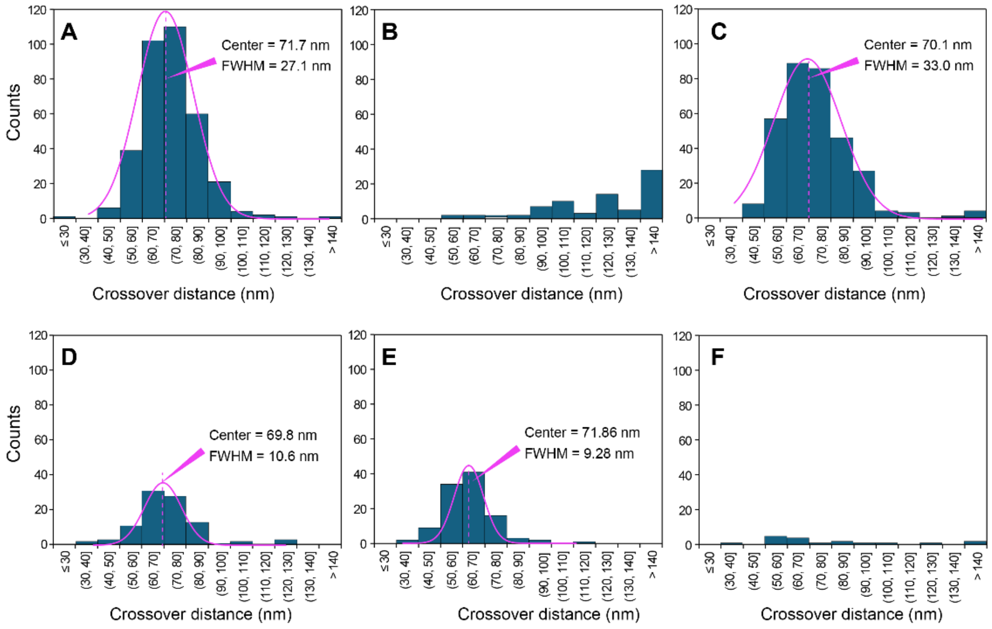


**Figure S4.** Histogram of crossover distance distribution for twisted α-syn fibrils formed in the presence of 2 (**A**; α-syn+2M), 4 (**B**; α-syn+4M), and 8 (**C**; α-syn+8M) mol% Aβ42 monomers or 2 (**D**; α-syn+2P), 4 (**E**; α-syn+4P), and 8 (**F**; α-syn+8P) mol% Aβ42 PFFs. Magenta lines represent Gaussian fits centered at 71.7, 70.1, 69.8, and 71.86 nm for A, C, D, and E, respectively. FWHM (full width at half maximum) describes the width of the crossover-distance distribution. The α-syn+8M (**C**) sample showed the broadest distribution (FWHM = 33.0 nm), while the α-syn+4P (**E**) sample showed the narrowest distribution (FWHM = 9.28 nm). Crossover distances were measured from 25 randomly selected TEM images for each sample.

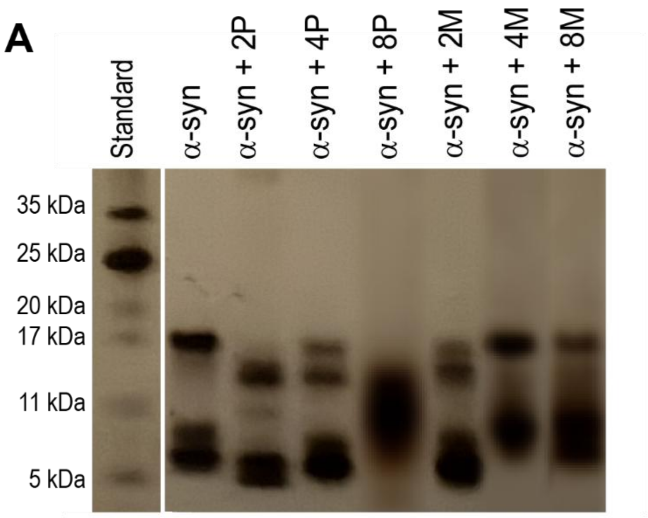


**Figure S5.** Proteinase K (PK) digestion profiles of α-syn fibrils formed in the absence or presence of Aβ42 monomers or PFFs. SDS-PAGE analysis of α-syn fibrils following proteinase K (PK) digestion.

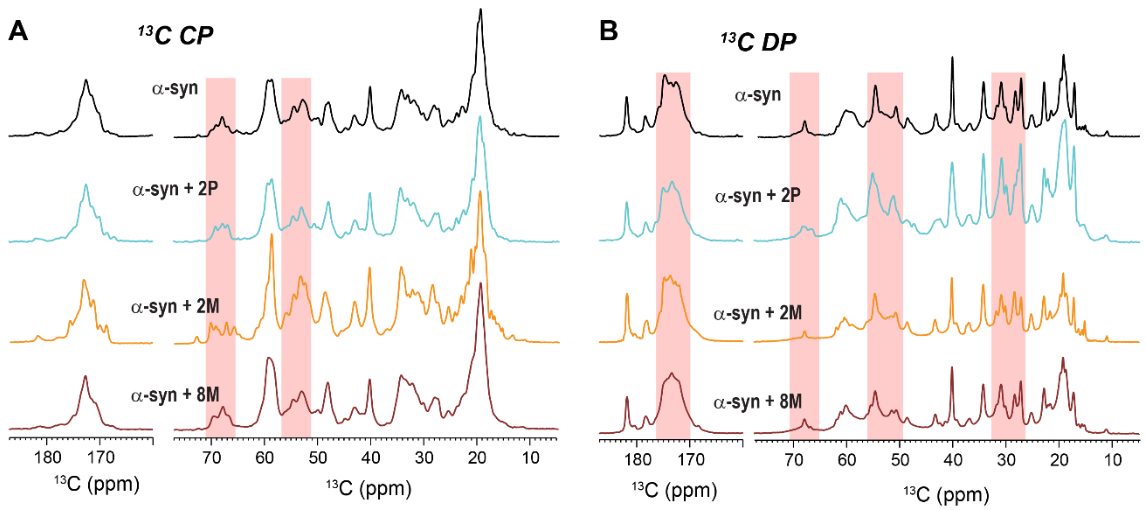


**Figure S6.** ssNMR 1D spectra. (**A**) 1D cross-polarization (CP) spectra and (**B**) 1D direct-polarization (DP) spectra of α-syn fibrils formed in the absence (α-syn, black) or presence of 2 mol% Aβ42 PFFs (α-syn+2P, cyan), 2 mol% Aβ42 monomers (α-syn+2M, orange), or 8 mol% Aβ42 monomers (α-syn+8M, dark red). Both CP and DP spectra show distinct spectral profiles among the samples, particularly in the Cα and carbonyl (CO) regions (red boxes), while the α-syn+2M sample showed a sharper and better-resolved CP spectrum.

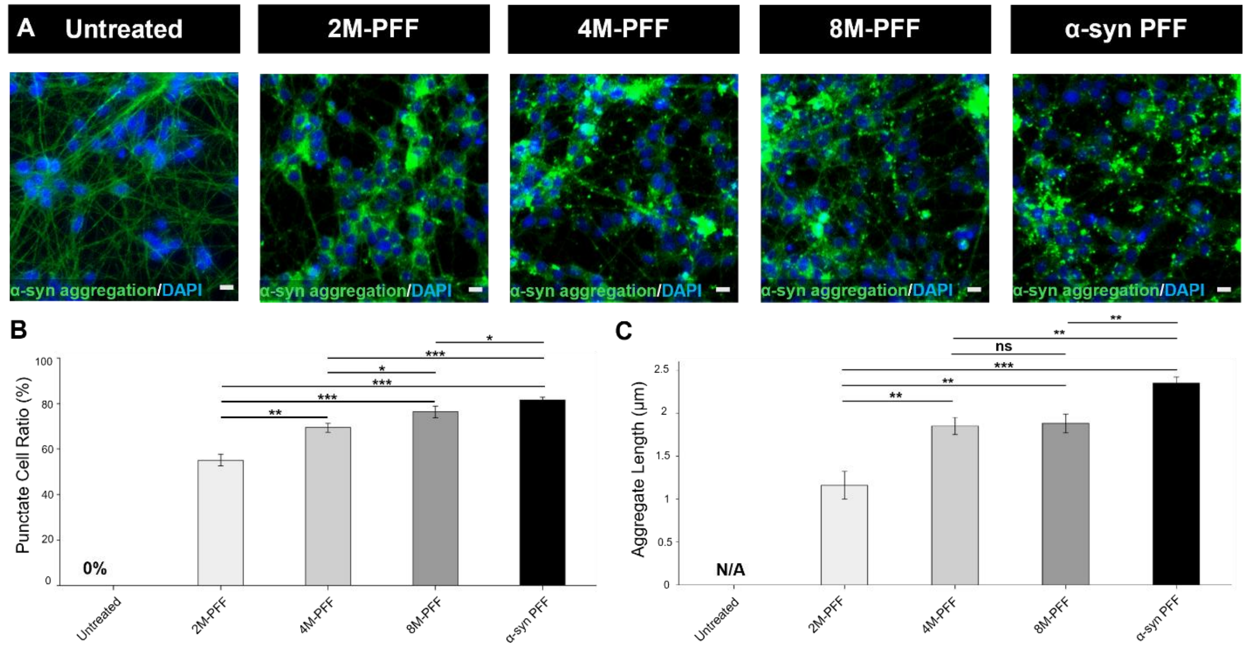


**Figure S7.** Intraneuronal α-syn aggregation induced by a reduced concentration (0.5 μM) of structurally distinct α-syn PFFs generated through heterotypic interactions with different concentrations of Aβ42 monomers. (A) Immunofluorescence staining for MJFR-14-6-4-2 (green) revealed intracellular α-syn aggregates in dopaminergic neuronal cells treated with heterotypic or homotypic α-syn PFFs, as well as in untreated controls. Scale bar, 10 μm. (B) Quantification of the puncta-positive cell ratio following PFF treatment compared with the untreated control. (C) Mean punctate aggregation length across heterotypic and homotypic α-syn PFF-treated groups and untreated controls. Data are presented as mean ± SD. *P < 0.05, **P < 0.01 and ***P < 0.001 (n = 3).

**Table S1.** Length distributions of Aβ42-modulated α-syn PFFs and homotypic α-syn PFFs.

| Sample | Mean PFF length (nm) | Median PFF length (nm) | SD (nm) | Minimum (nm) | Maximum (nm) | n |
|---|---|---|---|---|---|---|
| 2M-PFF | 76.4 | 74.1 | 19.6 | 43.5 | 138.9 | 80 |
| 4M-PFF | 78.5 | 77.4 | 19.4 | 37.7 | 122.3 | 80 |
| 8M-PFF | 83.6 | 78.4 | 21.8 | 42.6 | 138.9 | 80 |
| 2P-PFF | 80.5 | 79.3 | 20.1 | 39.4 | 137.2 | 80 |
| 4P-PFF | 77.1 | 76.6 | 22.7 | 36.4 | 126.8 | 80 |
| 8P-PFF | 80.3 | 79.4 | 19.4 | 43.3 | 130.0 | 80 |
| α-syn PFF | 81.9 | 82.2 | 18.1 | 47.2 | 130.6 | 80 |
